# Proteins are a source of glycans found in preparations of glycoRNA

**DOI:** 10.1101/2025.03.17.643711

**Authors:** Nathanael B. Kegel, Nurseda Yilmaz Demirel, Timo Glatter, Katharina Höfer, Andreas Kaufmann, Stefan Bauer

## Abstract

Recent discoveries show that RNA can be modified with sialylated glycans (termed glycoRNA), thus broadening our understanding of cellular glycosylation beyond traditional proteins and lipids. However, the pathway of RNA-glycosylation and its biological function remain elusive. Following the original glycoRNA isolation protocol, we also detect labelled glycans in small RNA preparations. However, glycosylated molecules showed resistance to treatment with RNase A/T1 but were sensitive to proteinase K digestion under denaturing conditions. Using liquid chromatography-mass spectrometry (LC-MS) based proteomics, we detect various proteins that co-purify with small but not large RNA preparations isolated from human or murine cells, including the glycosylated membrane protein LAMP1. Importantly, we further demonstrate that recombinant soluble LAMP1 can be purified following the glycoRNA isolation method. These findings suggest that glycoproteins co-purify with RNA using current glycoRNA purification protocols, thus representing a considerable source of glycans in samples of glycoRNA.

## Introduction

Glycosylation shapes the structure, interactions, and signaling properties of its targets^1,2^, which traditionally are considered to be proteins and lipids. Recently, Flynn et al. reported RNA as the newest member of the family of glycosylated biomolecules in living cells^3^. The group employed metabolic labelling with peracetylated N-azidoacetylmannosamine (Ac_4_ManNAz) to introduce a clickable azido-sialic acid to nascent N-glycans^4^, thus enabling the attachment of chemical reporters such as biotin to the labelled glycans by strain-promoted azide-alkyne cycloaddition (SPAAC)^5^. By using a streptavidin probe, labelled glycans were detected in RNA samples via Northern blotting. Signals displayed high molecular weight in the range of >10 kb and were lost upon disruption of the N-glycosylation machinery and upon RNase A/T1 treatment, yet remained unaffected by DNase I or proteinase K. Despite high apparent molecular weight, glycan incorporation was restricted to preparations of small RNAs of less than 200 bp. RNA sequencing of affinity-purified glyco-conjugates identified small non-coding RNAs, such as Y, U, small nucleolar, and transfer RNAs, as possible targets of RNA glycosylation. Interestingly, subcellular fractionation and subsequent RNA extraction demonstrated that glycan-modified RNAs are associated to the outside of cellular membranes, where they serve as ligands for sialic acid-binding lectins (Siglecs) 11 and 14. Together, these data show a cell surface-attached covalently linked RNA-glycan conjugate (hereafter: glycoRNA)^3^.

The discovery of glycoRNAs has raised questions regarding the nature of the RNA-glycan linkage, the underlying pathway by which sugar chains are attached to RNA, and the biological function^6–9^. Recently, leveraging a periodate-based method for the detection of glycans in native RNA, the non-canonical RNA base 3-(3-amino-3-carboxypropyl)uridine (acp^3^U) was reported as the N-glycan attachment site on RNA^10^. The functional aspects of glycoRNA are increasingly being uncovered with recent publications highlighting a role of glycoRNA in tumor progression and monocyte adhesion to endothelial cells^11^, neutrophil recruitment to inflammatory sites^12^, and involvement in the regulation of epithelial barrier function in the lung^13^. Moreover, tight interactions of glycoRNA with other components of the cell membrane, such as lipid rafts, cell-surface proteins, and heparan sulfate^11,14–16^ indicate the existence of complex microenvironments at the cell surface.

Even though the glycoRNA field is expanding with new discoveries, a current study has highlighted the sensitivity of glycoRNA detection to specific purification steps^17^. In line with this finding, we here show that the reported elimination of detectable glycans from RNA by RNase treatment depends on a post-digestion silica column purification step. We observed that the column binding of the labelled glycosylated molecules is impaired upon RNase treatment, and labelled glycans can be recovered based on the purification steps. Analysis of glycan-rich RNA by SDS-PAGE revealed the presence of a glycosylated molecule of approximately 100 kDa. Importantly, we show that this glyco-molecule is sensitive to proteinase K digestion under denaturing conditions. Mass spectrometry-based proteomics confirmed that several glycoproteins, among them LAMP1, co-purify with small RNA preparations and withstand the extraction strategy described in the glycoRNA protocol^3^. Our findings suggest that glycoproteins contribute to the level of detectable glycans in RNA preparations.

## Material & methods

### Cells

NIH3T3 cells (3T3 cells) were grown in DMEM (Anprotec, AC-LM0015) with 10% (vol/vol) fetal calf serum (FCS, Gibco), 2 mM L-glutamine (Gibco), 100 IU/mL penicillin-streptomycin (Gibco) at 37°C, 7.5% CO_2_. HeLa cells were cultivated in RPMI-1640 (Anprotec, AC-LM0051) with identical supplements at 37°C, 5% CO_2_. Cells were passaged twice per week in a 1:10 ratio in T25 plastic flasks.

### Metabolic labelling

N-glycans were metabolically labelled as described previously^3^. In brief, confluent 3T3 or HeLa cells were harvested by trypsin detachment. Cells were pelleted, resuspended in complete cell culture medium and counted using a Neubauer cell counting chamber. Typically, 4.9 × 10^6^ cells were seeded in a T175 cell culture flask. Labelling medium consisted of 25 mL DMEM or RPMI with supplements as stated above and 100 µM Ac_4_ManNAz (Jena Bioscience), 100 µM GalNAc (Sigma), and 10 µM D-Gal (Roth). Cells were incubated at 37°C, 7.5% CO_2_ for 40 hours.

### RNA extraction

After 40 hours of metabolic glycan labelling, culture medium was aspirated and adherent cells were washed once with PBS. TRIzol RNA extraction reagent (Invitrogen, 15596018), was added directly to the cells and incubated for 10 minutes at ambient temperature. For T175 flasks, 10 mL TRIzol was applied. Cells were homogenized by pipetting, transferred to a 15 mL reaction tube, and incubated for another 10 minutes at 37°C to increase lysis efficiency. To initiate phase separation, 0.2× volumes of chloroform (100%) were added and mixed by thorough shaking. After centrifugation at 4000 × g for 10 minutes, the aqeuous phase was transferred to a fresh reaction tube and mixed with 1.1× volumes of isopropanol (100%). RNA was precipitated at -20°C for one hour, then pelleted by centrifugation at 4000 × g, 4°C for two hours. The pellet was washed once with ethanol (80%), then air-dried using a laminar flow hood and lastly solubilized overnight in ultrapure water (UltraPure™, Invitrogen).

### Silica column purifications

Silica columns were used to de-salt RNA after TRIzol extraction and to remove metabolites and unconjugated click chemistry reagents following the adaptions described recently^3,18^. In brief, samples were mixed with two volumes of RNA Binding Buffer (Zymo Research, R1013-2) and briefly vortexed. An additional volume of isopropanol (100 %) was added (e.g. 100 µL of RNA + 200 µL RNA Binding Buffer + 300 µL isopropanol) and samples were mixed by vortexting and briefly incubated on ice. Samples were loaded to Zymo Spin™ silica columns and centrifuged for 30 seconds at 16,000 × g. For up to 70 µg of RNA, Zymo Spin™ IC columns (Zymo Research, C1004) were used, and IIICG columns (C1006) were used for RNA quantities of up to 350 µg^3^. Three separate washing steps were included: once with 400 µL RNA Prep Buffer (Zymo Research, R1060-2) and 30 seconds centrifugation, once with 700 µL ethanol (80%) and 30 seconds centrifugation, and once with 400 µL ethanol (80%) for 60 seconds to remove residual alcohol. RNA was eluted twice with ultrapure water. After the column purification, RNA concentration was quantified by NanoDrop® spectroscopy. Alternatively, when extracting lower amounts of RNA (e.g. from six-well plates), we found that precipitation from the aqeuous phase of TRIzol with two volumes of isopropanol (100%) and direct loading to the silica columns allowed for indentical results.

Separation of large (>200 nts) and small (<200 nts) RNAs was conducted according to the protocol described by Hemberger et al.^18^. In brief, total RNA was mixed with two volumes of an adjusted RNA Binding Buffer (equal parts of RNA Binding Buffer and 100% ethanol) and loaded to a silica column. After centrifugation, the large RNA fraction was retained in the column, and the flow-through, containing the small RNA fraction, was mixed with an additional volume of isopropanol (100%). The small RNA fraction was loaded to a separate Spin™ column. Purification proceeded as described above.

### Proteinase K treatments

For further RNA sample purification, RNA de-salted via silica column purification was treated with proteinase K as described previously^3^. In brief, proteinase K (PCR grade, Thermo Scientific, EO0491) was added directly to RNA in ultrapure water at a concentration of 1 µg proteinase K per 25 µg of RNA. The treatment was performed for 45 minutes at 37°C. Alternatively, treatment was conducted in denaturing Tris buffer (DTB) to provoke protein unfolding and enhance enzymatic activity^19^. DTB was prepared analogous to SDS-PAGE sample buffer^20^ containing 375 mM Tris-HCl (pH 6.8), 9% SDS, 10% 2-mercaptoethanol, 50% glycerol and 0.003% bromophenol blue. Typically, 25 µg of RNA in 40 µL of ultrapure water was supplemented with 1 µg of proteinase K, then 14 µL of ultrapure water or DTB was added (final buffer concentrations: 97.2 mM Tris-HCl, 2.3% SDS, 2.6% 2-mercaptoethanol, 13% glycerol). After 45 minutes of treatment, samples were filled to 100 µl with ultrapure water and buffers and metabolites were removed by another silica column clean-up as described above. All RNAs in this study were purified by these steps, except when specified otherwise.

To explore proteinase K activity against the glycosylated molecules in preparations of RNA over time, reaction tubes containing 2 µg of labelled, purified and clicked RNA (see below) in ultrapure water was supplemented with 1 µg of proteinase K per reaction tube and then transferred to a heating block at 37°C. Next, 0.4× volumes of DTB were added in intervals of 15 minutes, where the first tube received DTB after 0 minutes and the last tube received DTB 90 minutes after the first tube. Typically, 4 µL of DTB were added to 10 µL of the RNA-proteinase K mixture. Samples that received 4 µL of H_2_O instead of DTB were used as a control. After 90 minutes, 4 µL of DTB was applied to all samples that had not previously received DTB, while others received 4 µL of H_2_O to yield a total volume of 18 µL in all samples. Probes were then immediately and simultaneously boiled at 95°C for three minutes and subsequently analysed via SDS-PAGE and fluorescence in-gel detection (see below).

### Bioorthogonal click chemistry

Click reactions were performed following the procedure described by Flynn et al. (2021)^3^. In short, de-salted and proteinase K-treated RNA was subjected to copper-free click chemistry with 500 µM DBCO-biotin or -Alexa 647. Typically, 9 µL of RNA was mixed with 10 µL of df-GLB (final concentration: 50% formamide, 9 mM EDTA, and 0.0125% SDS)^3^ and 1 µL of DBCO-PEG_4_-biotin (Jena Bioscience) or DBCO-AF647 (Jena Bioscience). We found this reaction to be scalable without evident reduction of signal intensity, and results were reproducible for RNA inputs between 10 and 300 µg. Conjugation was performed at 55°C for 10 minutes. Afterwards, samples were filled to 100 µL with ultrapure water and excess click reagent was removed by another silica column clean-up as described above.

### Gel electrophoresis, RNA and glycan visualization

Glycan incorporation in RNA preparations was analyzed by fluorescence in-gel detection or by Northern blotting. For fluorescence in-gel detection, electrophoresis was performed in agarose gels with 1% agaorse (m/vol) and 1.5× MOPS buffer, submerged in 1× MOPS buffer (200 mM MOPS, 50 mM sodium acetate, 10 mM EDTA, adjusted to pH 7 with NaOH). Prior to loading, RNA samples were mixed with 5× denaturing loading buffer (50% formamide, 5.92% formaldehyde, 6.67% glycerol, 1× MOPS buffer) to yield at least 2× buffer concentration. Samples were denatured at 70°C for one minute and immediately placed on ice. Electrophoresis was conducted for 45 minutes at 105 V. Glycans were visualized directly in the agarose gel by fluorescence detection in the Alexa 647 channel. RNA was visualized by incubating the agarose gel in ethidium bromide solution (2 µg/mL) for 10 minutes at ambient temperature. RNA was detected by ultraviolet light using a GelStick Imager (Intas).

For glycan detection by Northern blotting, downward alkaline blotting^21^ was employed for transfer to a nitrocellulose membrane using a transfer buffer with 3 M NaCl and 8 mM NaOH. Transfer was performed for 150 minutes at ambient temperature. Afterwards, RNA was immobilized by UV irradiation (1.23 J/cm^2^) and the membrane was washed twice with PBS for removal of transfer buffer. The membrane was blocked overnight in blocking buffer (BB: 0.5% BSA in PBS with 0.1% Tween®-20). Streptavidin-HRP (Roche) was diluted 1:5,000 (vol/vol) in BB and used for detection of DBCO-biotin-conjugated labelled glycans. Signals were detected with Femto (Thermo Scientific, 34096) and a ChemiDoc MP imaging system. Afterwards, total RNA was stained on the membrane with methylene blue^22^. In brief, the nitrocellulose membrane was pre-incubated in 5 % acetic acid for 15 minutes at ambient temperature. Staining was conducted in 0.5 M sodium acetate (pH 5.2) and 0.04% methylene blue for 10-30 minutes at ambient temperature until clear ribosomal RNA bands were visible. The membrane was washed several times with de-salted water and then photographed using a ChemiDoc MP imager.

To probe for presence of proteins and investigate protein-like migration of glycan-bearing molecules in preparations of RNA, clicked samples of RNA were analyzed by sodium dodecylsulfate polyacrylamide gel electrophoresis (SDS-PAGE) following the procedure described by Laemmli (1970) using gels with 10 % polyacrylamide^20^. RNA samples were mixed with SDS-PAGE sample buffer containing SDS and 2-mercaptoethanol and were immediately boiled at 95°C for three minutes. Gels were run for at least two hours until the bromophenol blue front reached the lower gel end and glycans were detected by in-gel fluorescence as explained above. PageBlue Protein Staining Solution (Thermo Scientific, 24620) was used to stain for proteins.

### RNase A/T1 treatments

To reproduce recent data concerning RNase sensitivity of glycoRNA^3^, 20 µg of labelled, purified and clicked RNA was mixed with 1 µL RNase A/T1 cocktail (Thermo Scientific), containing 0.5 U RNase A and 20 U RNase T1, in RNase Buffer (20 mM Tris-HCl at pH 8.0, 100 mM KCl and 0.1 mM MgCl_2_). Samples were incubated for 60 minutes at 37°C. Afterwards, to compare glycan abundance and migration behaviour before and after silica column purification, samples were split in half, where one portion was put on ice and the other was subjected to a silica column purification as explained above. Samples were analyzed by agarose gel electrophoresis and fluorescence in-gel detection.

To investigate the importance of intact RNA for the precipitation of glycoproteins in samples of RNA, 16 µg of labelled, purified and clicked RNA was mixed and incubated with RNase A/T1 as stated above. Samples without RNase A/T1 were used as a control. An aliquot equivalent to 2 µg of RNA from each group was separated as an “input” control. Next, samples were split in three, where each reaction tube was mixed with two volumes of RNA Binding Buffer (Zymo). Importantly, RNA Binding Buffer contains guanidine salts, which inactivate the RNase. The first tube received another volume of isopropanol (100%). The second tube was spiked with 5 µg of intact RNA from 3T3 cells not exposed to Ac_4_ManNAz-labelling and then received another volume of isopropanol (100%). The third tube received two volumes of isopropanol (100%). Column purification proceeded as explained above. The input controls and eluates from the columns were then analysed via agarose gel electrophoresis.

### Amicon® concentration

To probe for impaired column binding of glycosylated molecules following RNase A/T1 treatment, the silica column’s first flow-through (see above) was collected and immediately loaded to an Amicon® Ultra centrifugal filter device with a 10 kDa molecular weight cut-off (Milipore, UFC201024) for concentration. The filter membrane was washed three times with 1 mL of ultrapure water and centrifuged at 2,800 × g for 15 minutes at 4°C. Concentrated sample was eluted by inversion of the filter device and centrifugation at 2,800 × g for one minute, which spins the sample out into a fresh reaction tube. The sample volume was further reduced by vacuum centrifugation. Analysis was conducted by fluorescence in-gel detection.

### LC-MS sample preparation and analysis

For sample preparation, total RNA was extracted from 3T3 or HeLa cells with TRIzol and was de-salted using silica columns as described above. A portion of this RNA was separated (total RNA). The rest received 1 µg proteinase K per 25 µg RNA, which was directly added to the purified RNA in ultrapure water. After incubation for 45 minutes at 37°C, large (>200 nts) and small (<200 nts) RNA fractions were separated as explained above. 100 µg of RNA from each sample group (total RNA, large RNA fraction, small RNA fraction) was digested with 1 µL PNGase F (NEB, P0704S) in 1× GlycoBuffer 2 (NEB) by overnight incubation at 37°C to remove N-glycans. This was followed by the addition of 1 µL RNase A/T1 cocktail (Thermo Fisher) at 37°C for 30 minutes. A short sonication step (30 seconds, 80% amplitude, 0.5 pulse) was performed to degrade remaining nucleic acids. 100 µL hot lysis buffer (100 mM ammonium bicarbonate, 2% SLS, 10 mM TCEP, 95°C) was added per 100 µL sample and samples were boiled for 10 minutes at 95°C. Next, 4 mM iodoacetamide was added to the samples and incubated for 30 minutes protected from light. 40 µL of Sera-Mag magnetic carboxylate-modified particles (Cytiva) and 40 µL of SpeedBead magnetic carboxylate-modified particles (Cytiva) were mixed, washed three times with MQ water and resuspended in 200 µL MQ water. 10 µL of this bead suspension was used per sample to capture proteins following the manufacturer’s instructions. On-bead tryptic digestion of proteins was performed by adding 1 µg of sequencing-grade trypsin (Promega) to the beads and overnight incubation at 30°C and 1200 rpm. Subsequently, the supernatant was collected and residual SLS was precipitated by adding 1.5% trifluoroacetic acid and subsequent centrifugation at 17,000 × g for 10 minutes at 4°C. The supernatant was separated and further desalted for mass spectrometric analysis using C18 solid phase columns (Chomabond C18 spin columns; Macherey-Nagel).

The peptides were dried, reconstituted in 0.1% TFA and then analyzed using liquid-chromatography-mass-spetrometry (LC/MS) carried out on an Exploris 480 instrument connected to an Vanquish Neo HPLC and a nanospray flex ion source (all Thermo Scientific). Peptide separation was performed on a reverse phase HPLC column (75 μm × 42 cm) packed in-house with C18 resin (2.4 μm; Dr. Maisch). The following separating gradient was used: 94% solvent A (0.15% formic acid) and 6% solvent B (99.85% acetonitrile, 0.15% formic acid) to 35% solvent B over 30 minutes at a flow rate of 300 nl/min. Peptides were ionized at a spray voltage of 2.3 kV, ion transfer tube temperature set at 275°C, 445.12003 m/z was used as internal calibrant. The data acquisition mode was set to obtain one high resolution MS scan at a resolution of 60,000 full width at half maximum (at m/z 200) followed by MS/MS scans of the most intense ions within one second (cycle 1 s). To increase the efficiency of MS/MS attempts, the charged state screening modus was enabled to exclude unassigned and singly charged ions. The dynamic exclusion duration was set to 14 seconds. The ion accumulation time was set to 50 ms (MS) and 50 ms at 17,500 resolution (MS/MS). The automatic gain control (AGC) was set to 3 × 10^6^ for MS survey scan and 2 × 10^5^ for MS/MS scans. The quadrupole isolation was 1.5 m/z, collision was induced with an HCD collision energy of 27%. MS raw data was then analyzed with MaxQuant^23^, and a *Mus musculus* and *Homo sapiens* uniprot database, respectively (Supplementary Data File). MaxQuant was executed in standard settings without “match between runs” option. The search criteria were set as follows: full tryptic specificity was required (cleavage after lysine or arginine residues); two missed cleavages were allowed; carbamidomethylation (C) was set as fixed modification; oxidation (M), deamidation (N,Q) as variable modification.

### Soluble human LAMP1

For expression of soluble human LAMP1, the genetic information for LAMP1 aa 29 to 365 (accession NP_005552.3) was amplified from HeLa cDNA with primers sol_hs_LAMP1_f_AfeI: 5’ATATAAGCGCTGCAATGTTTATGGTGAAAAATG and sol_hs_LAMP1_rev_BamHI: 5’ATATGGATCCTCAGTTCTCGTCCAGCAGACAC. The fragment was purified and digested with AfeI and BamHI and cloned into an equivalently treated pcDNA3.1(-) vector originally containing solBDCA2 (accession number OQ817991). The generated construct consists of an N-terminal HLA-A2 signal peptide (MAVMAPRTLLLLLSGALALTQTWAGSHS; P04439.2, aa 1-28), a Twin-Strep-tag (WSHPQFEKGGGSGGGSGGSAWSHPQFEKSA) (IBA) followed by LAMP1 aa 29 to 365. The sequence of the LAMP1 fragment was verified by Sanger sequencing. For protein expression, HEK293 cells were transfected using 0.1 µg of the corresponding plasmid and Lipofectamin™ 2000 in a 96-well format according to the manufacturer’s protocol. Transfected cells were selected and cloned using 0.7 µg/mL G418 (Thermo Fisher Scientific). The purification of secreted proteins from the supernatant was carried out using Strep-Tactin cartridge (IBA) according to the manufacturer’s instructions. Proteins were concentrated after purification with centrifugal filters (10 kDa cut-off, Millipore) and washed to remove excess of biotin. Final protein concentration was measured using a BCA assay kit (Thermo Fisher Scientific). The proteins were analyzed by SDS-PAGE and subsequent Coomassie staining (see above) or Western blot analysis (see below).

### Sol-hLAMP1 purification studies

Sol-hLAMP1 was used as a tool to investigate the possibility of glycoprotein co-purification following the published glycoRNA isolation method^3^. To test its phase separation behaviour, 2 µg of purified sol-hLAMP1 in 20 µL of ultrapure water was mixed with 1 mL of TRIzol by rigorous pipetting and then incubated and mixed with chloroform as explained above. The aqeuous phase was carefully extracted and mixed with two volumes of isopropanol and then purified using a silica column. To control direct binding to silica columns, 2 µg of sol-hLAMP1 in 100 µL of ultrapure water was mixed with two volumes of RNA Binding Buffer (Zymo) by vortexing. Next, one, two or three volumes of ethanol or isopropanol (100%) were added and samples were vortexed and briefly incubated on ice. The silica column purification continued as explained above. Yields of sol-hLAMP1 were assessed by Western blotting (see below).

To investigate the influence of intact RNA for adherence of sol-hLAMP1 to silica columns, an experiment identical to the one performed with clicked RNA (see section “RNase A/T1 treatments”) was conducted using 2 µg of sol-hLAMP1 in 20 µl for each sample. In brief, sol-hLAMP1 was left untreated, treated with 1 µl RNase A/T1 in RNase buffer as explained above, spiked with 5 µg of RNA from 3T3 cells not exposed to Ac_4_ManNAz labelling, or spiked with RNA and additional RNase A/T1. All samples were incubated for 45 minutes at 37°C. Aliquots equivalent to 1 µg sol-hLAMP1 were then separated as “input” controls. The rest was mixed with two volumes of RNA Binding Buffer (Zymo) and another volume of isopropanol (100%). Two additional controls were used: one sample was mixed with RNase A/T1 and then with RNA Binding Buffer (Zymo) to inactivate the RNase, then received 5 µg of RNA; another sample was mixed with two instead of one volume of isopropanol. All samples were then purified using Zymo Spin™ IC columns as per the protocol explained above.

### Western blot analysis of sol-hLAMP1 and endogenous LAMP1

Following analysis by SDS-PAGE as explained above, proteins were transferred to polyvinylidene fluoride (PVDF) by semi-dry Western blotting. PVDF membranes were blocked overnight in TBS-T containing 5% bovine serum albumin (BSA). Sol-hLAMP1 was detected using Strep-Tactin HRP conjugate (IBA) diluted 1:40,000 (vol/vol) in TBS-T with 5% BSA. Endogenous human LAMP1 was detected using rabbit anti-LAMP1 mAb (XP® D2D11, #9091, Cell Signaling Technology) diluted 1:1,000 (vol/vol) in TBS-T with 5% skim milk powder and peroxidase-conjugated goat anti-rabbit IgG (Jackson Immuno Research, 111-035-045) diluted 1:15,000 in TBS-T with 5% skim milk powder as secondary antibody. Actin staining served as a loading control using mouse monoclonal anti-beta-actin antibody (Sigma, A2228) diluted 1:10,000 (vol/vol) in TBS-T with 5% skim milk powder and peroxidase-conjugated anti-mouse IgG (Jackson Immuno Research, 115-035-062) as secondary antibody. Blots were washed three times with TBS-T before and after secondary antibodies were administred. Signals were detected using SuperSignal™ West Dura (Thermo Scientific, 34075) or Femto (Thermo Scientific, 34096).

## Results

### Labelled glycans are unaffected by RNase A/T1 treatment

Glycan modification of cellular RNA was investigated following the procedure described by Flynn et al. (2021)^3^. The workflow comprises metabolic labelling of cells with Ac_4_ManNAz, RNA extraction with TRIzol^24^, subsequent RNA purification with silica columns, proteinase K treatment, and a second silica column purification. To visualize labelled glycans, RNA samples were subjected to copper-free click chemistry with DBCO-biotin^3^. Large (>200 nts) and small (<200 nts) RNA fractions were separated and subsequently analysed by Northern blotting (Figure 1a). We observed labelled glycans in small but not large RNA fractions extracted from HeLa and NIH3T3 cells (3T3 cells) (Figure 1b) in agreement with previous reports^3,10–12^. Although restricted to the small RNA fraction, glycan-modified molecules displayed high apparent molecular weight in the range of 3-5 kb.

**Figure 1:**
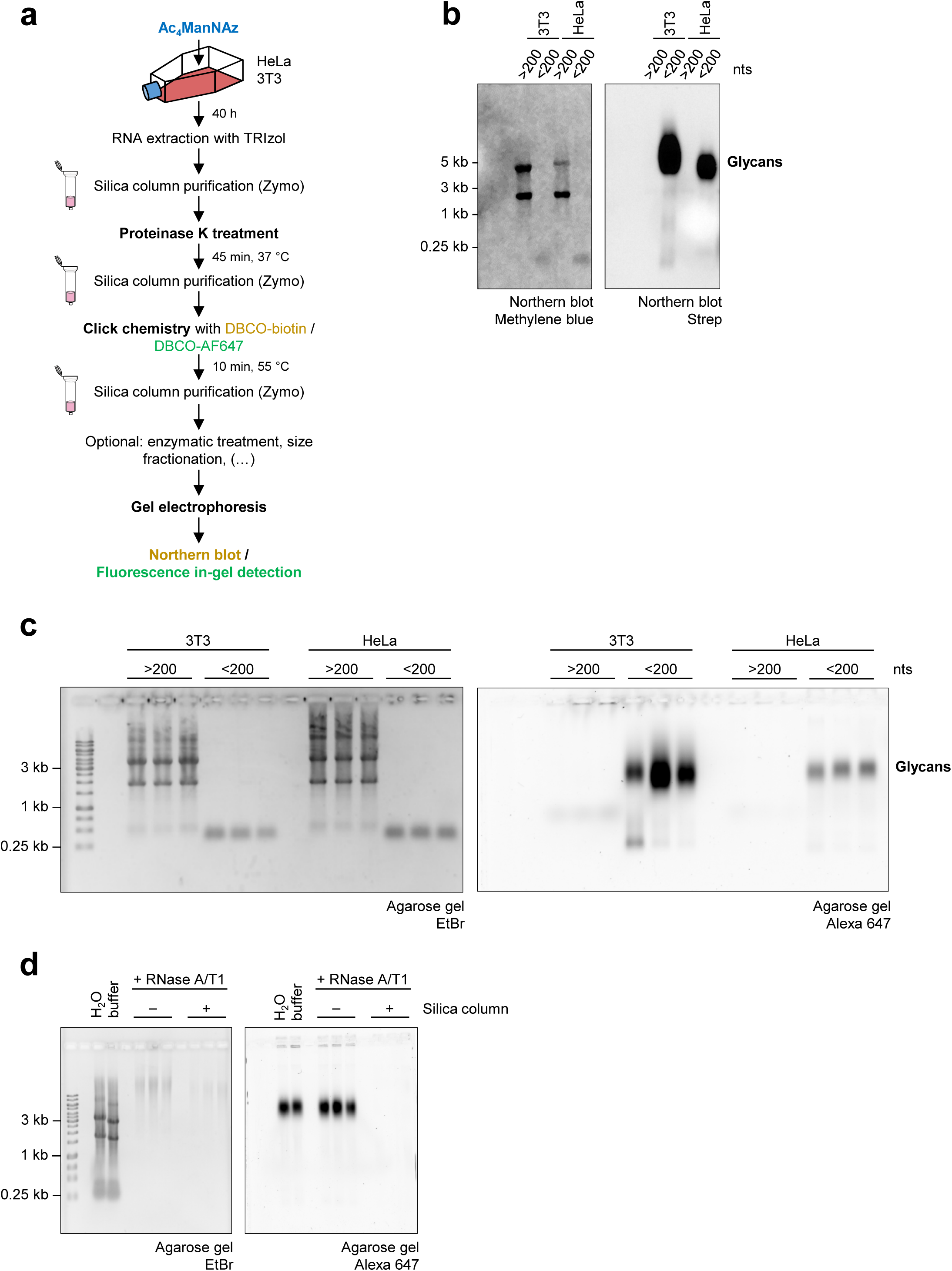
Preparations of glycoRNA contain RNase-resistant glyco-molecules. (a) Schematic representation of the glycoRNA labelling and purification strategy, following the procedure described by Flynn et al. (2021)^3^. (b) Northern blot of RNA extracted from metabolically labelled 3T3 and HeLa cells. RNA was purified as outlined in (a) and conjugated to DBCO-biotin. Methylene blue was used to detect RNA on the nitrocellulose membrane. A streptavidin-peroxidase probe was used to detect labelled glycans conjugated to DBCO-biotin. (c) In-gel fluorescence detection of labelled glycans in preparations of RNA extracted from metabolically labelled 3T3 and HeLa cells. RNA was purified as in (a) and conjugated to DBCO-AF647. Following gel electrophoresis, labelled glycans were visualized directly in the agarose gel. Ethidium bromide (EtBr) staining serves as a loading control. Biological triplicates are shown. (d) RNase A/T1 treatment of RNA extracted from metabolically labelled HeLa cells. The first two lanes show undigested controls in ultrapure water or in digestion buffer. After incubation with RNase A/T1, samples were split. One half was left untreated and the other was purified using silica columns. Three replicates of individual digestions are shown.

For a fast and linear workflow that allows for direct detection of labelled glycans in agarose gels, we used copper-free click chemistry to conjugate the fluorophore AF647 to labelled glycans in RNA extracted from metabolically labelled cells. Glycans detected by this in-gel fluorescence detection method displayed high apparent molecular weight and restriction to preparations of small RNA (Figure 1c), analogous to glycans detected by Northern blotting (Figure 1b), and all subsequent analyses were conducted following this method. To investigate if the glycans detected in RNA preparations are linked to RNA, we subjected DBCO-AF647-clicked RNA from Ac_4_ManNAz-labelled HeLa cells to RNase A/T1 treatment. Labelled glycans were lost after RNase A/T1 treatment and subsequent silica column purification (Figure 1d) as described by Flynn et al.^3^. However, when RNase A/T1-treated samples were analysed without post-digestion purification with silica columns, we observed that the migration and integrity of labelled glycans were unaltered (Figure 1d). This observation implied that glycoRNA samples contained not only glycoRNA but also additional labelled glycosylated molecules that were co-purified under the conditions used for glycoRNA isolation.

### Highly stable glycoproteins are present in preparations of glycoRNA

To explore if proteins were present in glycoRNA samples, we analysed small RNA fractions from Ac_4_ManNAz-labelled cells (Figure 1c) by SDS-PAGE (Figure 2a). Since the samples were digested with proteinase K to eliminate protein contamination^3^, we did not detect protein bands by Coomassie staining as expected. By contrast, we observed several bands with the most prominent signal at approximately 100 kDa by fluorescence in-gel detection (Figure 2a), possibly implying an insufficient sensitivity of the Coomassie staining. In the original protocol for glycoRNA isolation, proteinase K is directly added to purified RNA in ultrapure water^3^. We speculated that under these conditions some proteins might be protected from cleavage due to inaccessible cleavage sites. Therefore, we treated DBCO-AF647-conjugated RNA extracted from metabolically labelled cells with proteinase K in a denaturing Tris buffer (DTB), containing SDS and 2-mercaptoethanol. We performed proteinase K treatment under denaturing and reducing conditions to enable target protein unfolding and generate access to potentially hidden cleavage sites^19^. Strikingly, proteinase K treatment in DTB led to degradation of the glycosylated molecule at 100 kDa in a time-dependent manner (Figure 2b and Supplementary Figure 1a). No digestion was observed in the absence of proteinase K or when the treatment was conducted in water. Based on these findings, we hypothesized that using the original protocol^3^, in addition to glycoRNAs also glycoproteins might be enriched and co-purified with the RNA. To test this hypothesis, we isolated RNA from metabolically labelled cells with TRIzol and silica column purification. Purified RNA was digested with proteinase K in DTB or in ultrapure water. Subsequently, metabolites and buffers were removed by a silica column purification step and samples were clicked to DBCO-AF647 to visualize labelled glycans (Figure 2c). Significantly, digestion with proteinase K in DTB but not in water strongly reduced the level of detectable glycans without affecting RNA integrity (Figure 2d and Supplementary Figure 1b). This observation highlights that some glycoproteins withstand the original glycoRNA isolation strategy and may represent a source of glycan-modified biomolecules in RNA preparations.

**Figure 2:**
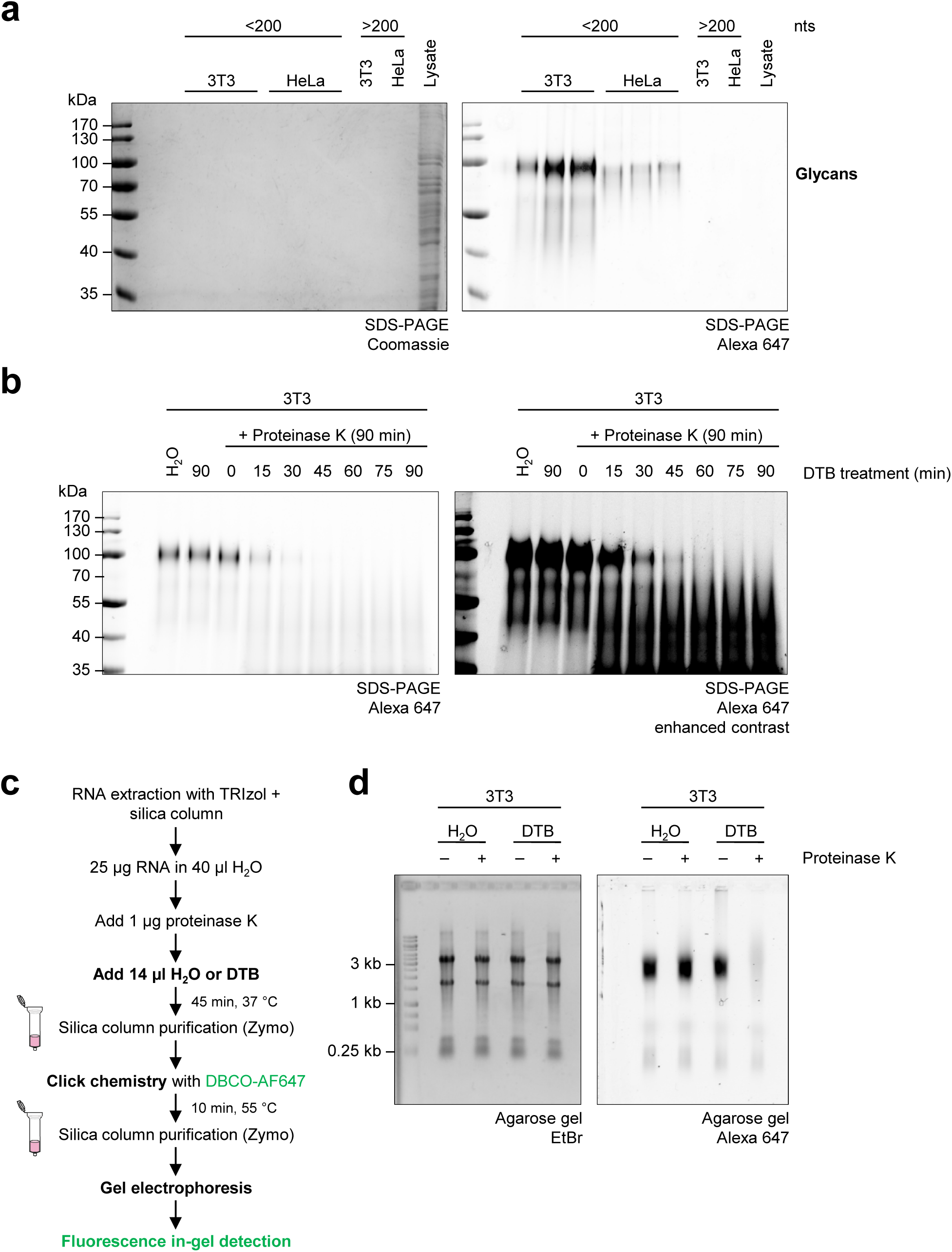
Proteinase K treatment in a denaturing buffer digests the glycosylated molecule. (a) SDS-PAGE of labelled RNAs from Figure 1c. Coomassie staining (left panel) was used to detect presence of proteins. Cell lysate obtained from native 3T3 cells serves as a positive control. Labelled glycans were visualized by fluorescence in-gel detection (right panel). Two samples of “large” RNA (>200 nts) fraction are shown as negative controls. (b) Time-course proteinase K digestion in denaturing Tris buffer (DTB). 1 µg of proteinase K was directly added to 2 µg of purified RNA in ultrapure water and incubated at 37°C for 90 minutes. DTB was added to the RNA-proteinase K mixture in intervals of 15 minutes as indicated. A representative gel from two independent experiments (n = 2) performed in two cell lines is shown (N = 4). (c) Schematic representation of a modified proteinase K treatment for glycoRNA purification (see Figure 1a) using DTB for denaturing and reducing reaction conditions. (d) In-gel fluorescence detection of labelled glycans in preparations of RNA extracted from metabolically labelled 3T3 cells, which were subjected to the conventional non-denaturing proteinase K treatment (see Figure 1a) or to the modified proteinase K treatment in DTB (see Figure 2c). Samples without proteinase K in ultrapure water or DTB serve as controls. A representative gel from two independent experiments (n = 2) in two cell lines is shown (N = 4).

In light of these findings, we revisited the observation that a post-digestion silica column purification step led to the loss of labelled glycans after RNase treatment (Figure 1d). We speculated that RNase A/T1 treatment might affect glycoprotein adherence to silica columns in an indirect fashion. Therefore, we analysed the flow-through of RNase A/T1-digested HeLa RNA subjected to column purification (Figure 3a). Importantly, after concentration using Amicon® Ultra columns with a 10 kDa cut-off filter, glycan-bearing molecules were retrieved from the flow-through showing no signs of degradation when analysed by agarose gel electrophoresis or SDS-PAGE (Figure 3b). These results demonstrated that the combination of RNase A/T1 treatment and silica column purification does not induce degradation but rather impairs the column binding ability of glycoproteins that act as contaminants in preparations of glycoRNA.

**Figure 3:**
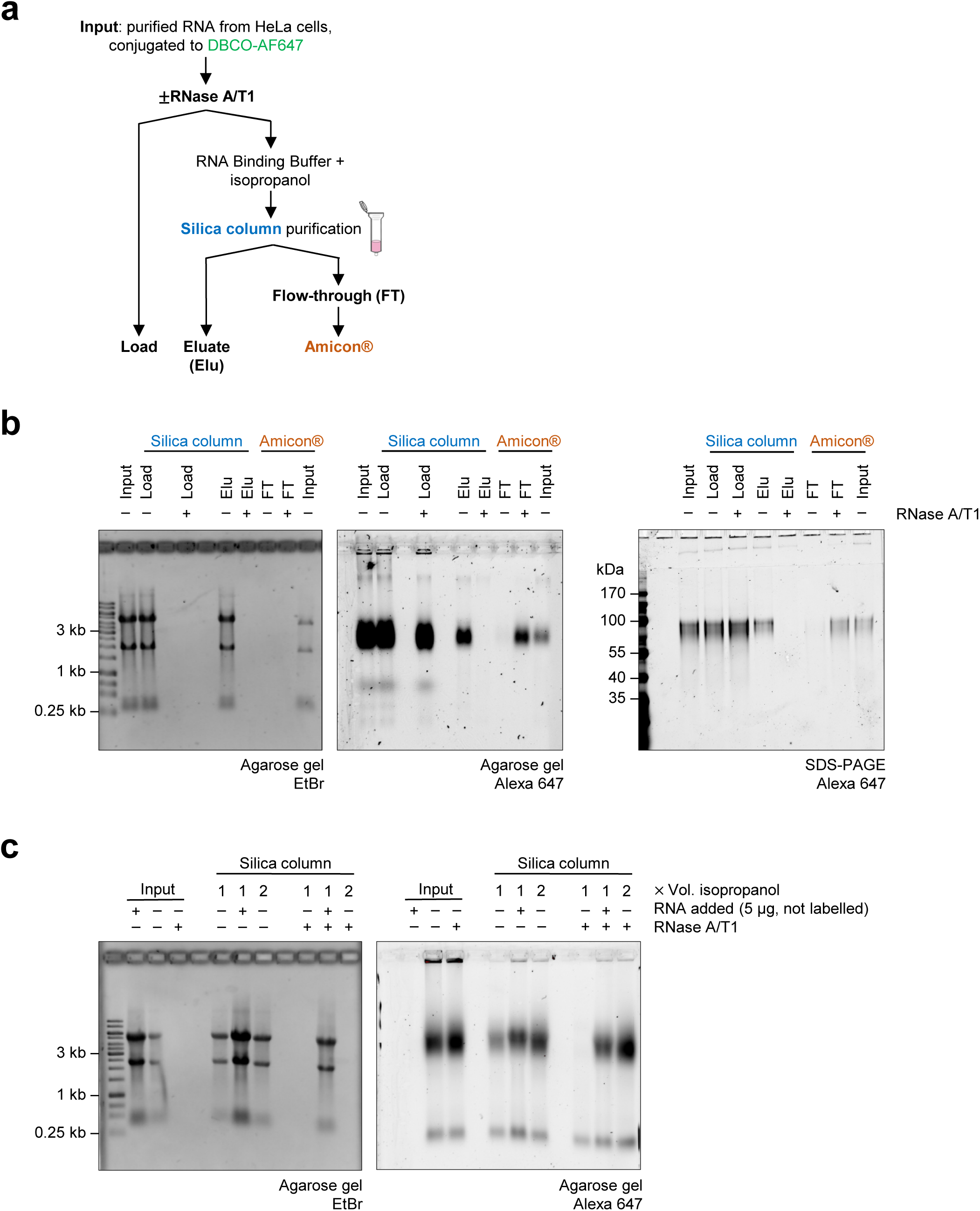
Loss of RNA impairs glycoprotein adherence to silica columns. (a) Schematic representation of a purification strategy to rescue glycosylated molecules from the flow-through of silica columns used for the purification of RNase-treated RNA samples. The flow-through of the first silica column loading step (see Material & Methods) is saved and transferred to a Amicon® concentrator column. (b) In-gel fluorescence detection of Ac_4_ManNAz-labelled glycans rescued from the flow-through of RNase-treated HeLa RNA. Lanes show an input control (“Input”), samples incubated with or without RNase A/T1 before column loading (“Load”), after elution (“Elu”), and the Amicon®-concentrated flow-through (“FT”). An untreated RNA (“Input”) was subjected to Amicon® concentration as a control. Identical samples were analyzed by agarose gel electrophoresis (left and middle panel) and SDS-PAGE (right panel). A representative gel from three independent experiments is shown (n = 3). (c) In-gel detection of labelled glycans obtained from silica column purifications in the presence of intact RNA or increased levels of isopropanol. RNA extracted from metabolically labelled 3T3 cells was incubated with or without RNase A/T1. After digestion, samples were split in three and then subjected to a silica column purification with three different conditions during the precipitation step (see Material & Methods): one tube received one volume of isoproanol, the second received 5 µg of intact RNA and one volume of isopropanol and the third received two volumes of isopropanol. Samples were purified in parallel and analyzed via agarose gel electrophoresis. Samples before the silica column run are shown as controls (“Input”). A representative gel of two independent experiments is shown (n = 2).

As Kim et al. (2024) recently described that intact RNA and increased concentrations of alcohol benefit the co-purification of N-glycosylated molecules when isolating glycoRNA^17^, we spiked RNase A/T1-treated RNA from metabolically labelled cells with intact RNA from cells not exposed to Ac_4_ManNAz-labelling or added an additional volume of isopropanol to the silica column’s precipitation step (see Material & Methods). Importantly, loss of labelled glycans after RNase A/T1 digestion and silica column purification could be prevented by both the presence of intact RNA and increased concentrations of isopropanol (Figure 3c). In summary, our observations suggest that highly stable glycoproteins withstand the purification strategy for glycoRNA. Their co-purification is mediated by the presence of intact RNA or high concentrations of alcohol during the column purification step.

### Identification of glycoprotein enrichment in small RNA preparations

To determine which proteins are co-purified with glycoRNAs, we developed a tandem liquid chromatography-mass spectrometry (LC-MS/MS) proteomics workflow. RNA from 3T3 and HeLa cells was isolated and de-salted using TRIzol and silica columns as described previously^3^. The RNA was left untreated (total RNA) or was treated with proteinase K, followed by large and small RNA separation^18^ (Supplementary Figure 2a, b). Samples of all three groups (total RNA, large RNA fraction, small RNA fraction) were digested with PNGase F and RNase A/T1 to remove N-glycans and RNA, respectively, and then subjected to the proteomics pipeline. To ensure confident protein identification, only proteins with an average unique peptide count of ≥2 across small RNA fraction triplicates were taken into consideration (Supplementary Table 1, 2). Principal component analysis (PCA) of sample label-free quantification (LFQ) intensity values revealed the close clustering of replicates from the same sample set for both cell lines (Supplementary Figure 2c, d). The heatmap illustrating log_2_-transformed LFQ intensities across large RNA fraction, small RNA fraction, and total RNA samples isolated from 3T3 cells (Figure 4a) revealed distinct protein detection profiles. As expected, proteins identified in the small RNA fraction samples were also detected in the total RNA samples, with comparable intensities. On the other hand, the majority of proteins detected in the small RNA fraction and total RNA samples were absent in the large RNA fraction samples. The heatmap of log_2_-transformed LFQ protein intensities in large RNA fraction, small RNA fraction, and total RNA samples isolated from HeLa cells (Figure 4b) exhibits a similar protein detection profile to the samples of 3T3 cells. Importantly, 10 proteins were commonly detected in the small RNA fractions of both HeLa and 3T3 cells, namely LAMP1, LAMP2, ITB1, 4F2, CALR, CD44, CD63, NUCKS, RL40, and BASI (Supplementary Figure 2e, f). It is important to mention that RL40, CD63, and NUCKS were also detected in the large RNA fraction samples isolated from 3T3 cells. Similarly, RL40, NUCKS, 4F2, and CD63 were present in the large RNA fraction samples isolated from HeLa cells. Taken together, these data suggest that specific proteins co-purify within preparations of proteinase K-treated small RNA fractions.

**Figure 4:**
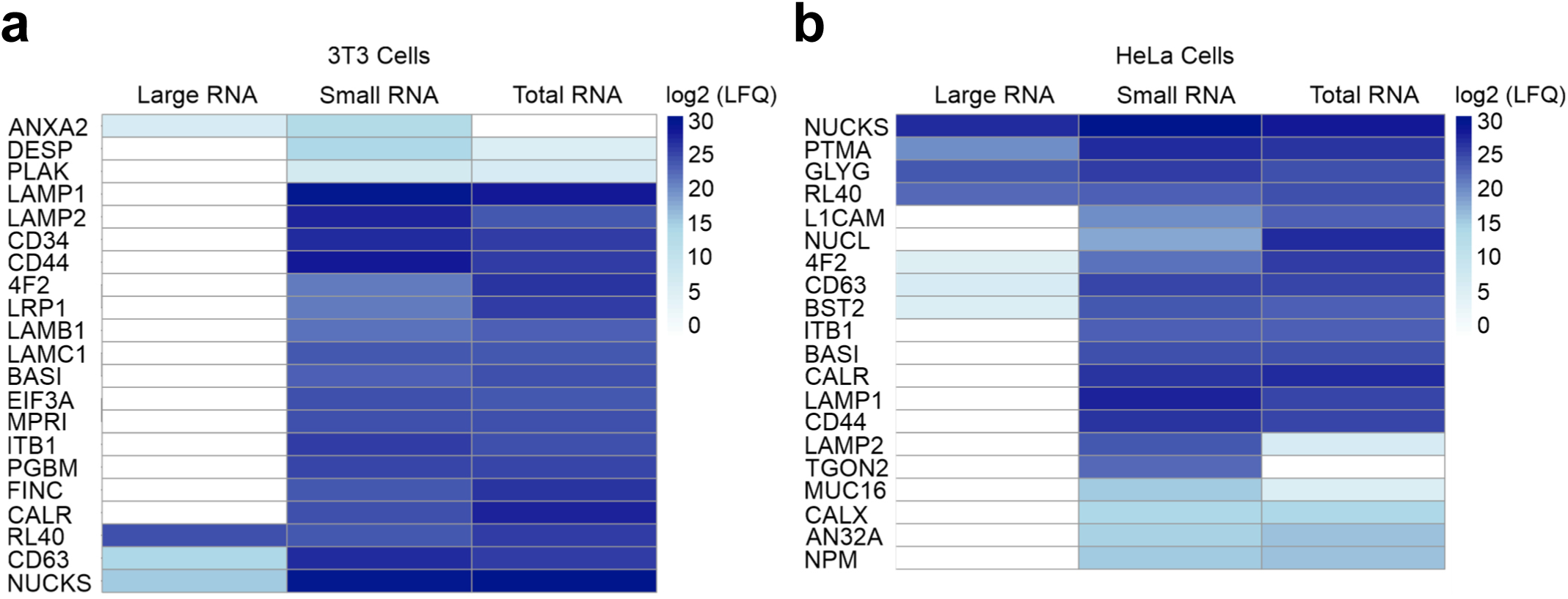
**Identification of proteins that co-purify with preparations of total, small, and large RNA fractions.** The heatmaps display the log_2_-transformed label-free quantification (LFQ) intensities of identified proteins across large RNA fraction, small RNA fraction, and total RNA samples from 3T3 cells (a) or HeLa cells (b), each analyzed in triplicates. The color scale represents intensity values, with darker shades indicating higher values.

### Recombinant LAMP1 can be purified using TRIzol and silica columns

Given the identification of LAMP1 in the proteomics data (Figure 4), and its description as a glycosylated membrane protein^25–27^, we studied the co-purification of LAMP1 under the conditions of the glycoRNA purification protocol^3^. We generated a recombinant Twin-StrepTag-containing soluble human LAMP1 (sol-hLAMP1), which showed similar glycosylation compared to endogenous LAMP1 as indicated by the mass increase to more than 100 kDa (Figure 5a). First, we investigated the phase separation of sol-hLAMP1 during TRIzol extraction and its ability to bind the silica columns used for RNA purification. To this end, sol-hLAMP1 was mixed with TRIzol, the aqueous phase was separated and two volumes of isopropanol were added. Alternatively, sol-hLAMP1 was directly loaded to the silica columns in the presence of one, two or three volumes of alcohol to test for the influence of increased alcohol concentrations for sol-hLAMP1 recovery. Remarkably, sol-hLAMP1 was readily detectable after extraction with TRIzol (Figure 5b), indicating that sol-hLAMP1 showed hydrophilic phase separation behaviour. Interestingly, one volume of either ethanol or isopropanol did not support its recovery. By contrast, two and three volumes of ethanol or isopropanol allowed for efficient sol-hLAMP1 purification.

**Figure 5:**
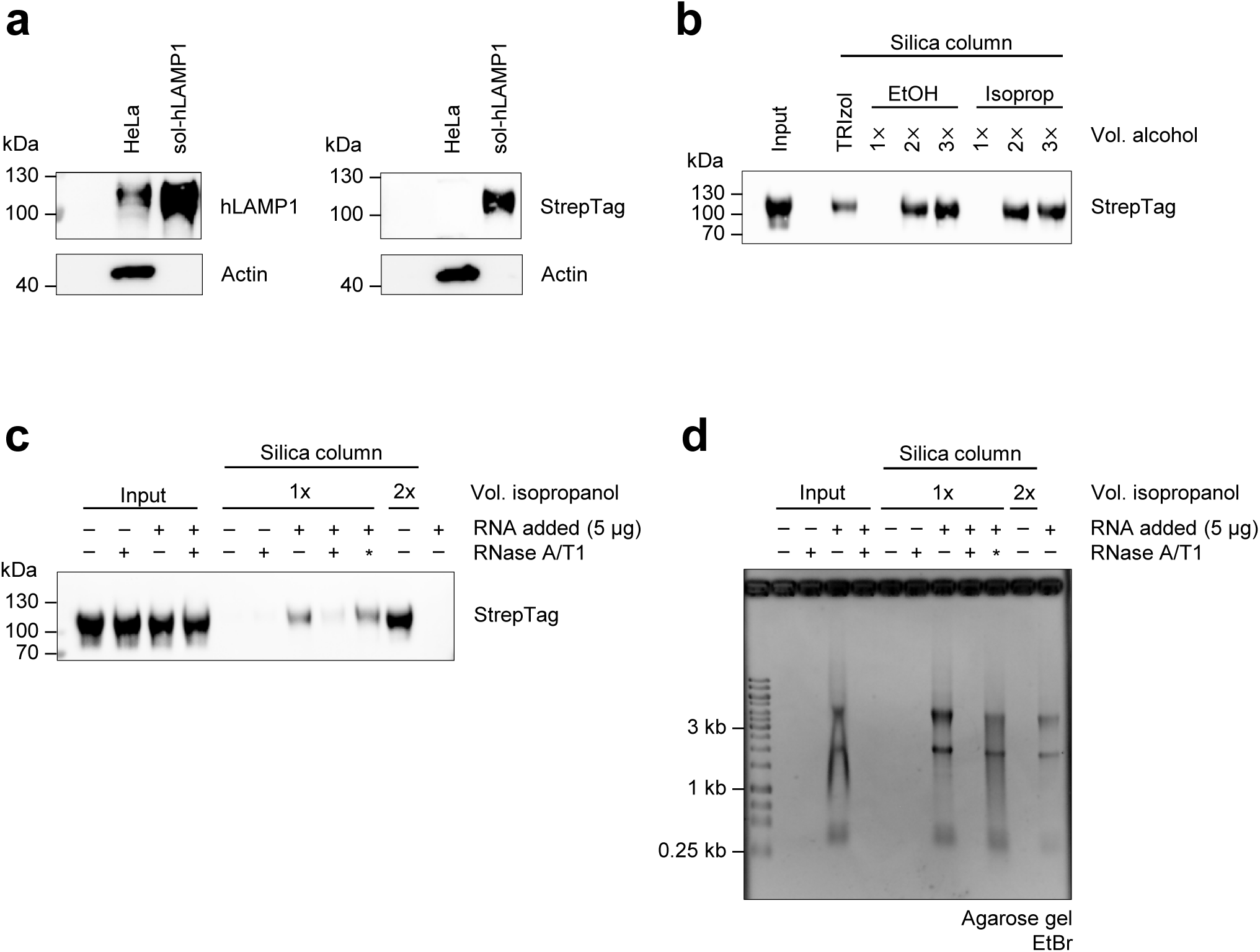
sol-hLAMP1 can be extracted from TRIzol and binds to silica columns. (a) Western blot analysis of the purified sol-hLAMP1 contruct compared to endogenous LAMP1 obtained from a HeLa cell lysate. (b) Western blot analysis of sol-hLAMP1 extracted via TRIzol or bound directly to a silica column. For direct column loading, samples were mixed with the silica column’s RNA Binding Buffer (Zymo) and additional volumes of ethanol or isopropanol as indicated. In case of the TRIzol treatment, sol-hLAMP1 was extracted from the aqeuous phase and mixed with two volumes of isopropanol. All samples were then purified using silica columns according to the manufacturer’s protocol and analysed by Western blotting. A representative blot of two independent experiments (n = 2) is shown. (c) Western blot analysis of sol-hLAMP1 obtained from silica column purifications in the presence of intact RNA or increased levels of isopropanol, analogous to Figure 3c. Sol-hLAMP1 was mixed with RNA and/or RNase A/T1 and incubated at 37°C for 45 minutes. Aliquots were separated (“Input”) and the rest was mixed with RNA Binding Buffer (Zymo) and isopropanol as indicated. The asterisk indicates inactivation of RNase A/T1 by guanidine salts before adding RNA. Silica column purification was carried out following the manfacturer’s protocol. A representative blot of two independent experiments (n = 2) is shown. (d) Agarose gel with EtBr staining of samples shown in (c).

To investigate if the presence of RNA promotes the purification of sol-hLAMP1, similar to experiments conducted by Kim et al. (2024)^17^ (and Figure 3c), we spiked sol-hLAMP1 with native RNA and performed a silica column purification using one volume of isopropanol for precipitation. RNase A/T1 treatment was used as a control. Before the column purification, neither RNase A/T1 treatment nor the addition of RNA affected the detectability of sol-hLAMP1 (Figure 5c, d). After the column purification, untreated or RNase A/T1-treated sol-hLAMP1 was no longer detectable. By contrast, sol-hLAMP1 could be recovered upon addition of RNA, which could be reverted by RNase A/T1 treatment. Inactivation of RNase A/T1 rescued sol-hLAMP1. As expected, the addition of two volumes of isopropanol allowed for the most efficient recovery. Taken together, our results indicate that sol-hLAMP1 as a representative glycoprotein has the physicochemical properties that allow co-purification during extraction of glycoRNA^3^. In addition, the presence of intact RNA or high alcohol concentrations are factors that enhance the co-purification of glycoproteins via silica columns.

## Discussion

Metabolic labelling is one of the core methods that enabled the identification and analysis of glycoproteins^4,28,29^. Flynn et al. used metabolic labelling with bioorthogonal sugar analogues to demonstrate that also RNA can be modified by glycans^3^, creating the emerging field of glycoRNA^3,11–16^. One key property of glycoRNA is its sensitivity towards RNase treatment as reported before^3^. However, we observed that glycoRNA preparations contain additional RNase-resistant glyco-molecules, whose loss after RNase treatment depends on a subsequent silica column purification step. In line with findings by Kim et al., we demonstrate that these glyco-molecules can adhere to silica columns in an RNA- and alcohol concentration-dependent manner^17^. Combining the observations that proteinase K digestion under denaturing conditions led to the degradation of these molecules, and the identification of 20 and 21 proteins in small RNA fractions extracted from 3T3 and HeLa cells, respectively, we conclude that proteins may co-purify with RNA under the original glycoRNA purification protocol^3^.

The presence of proteins in samples of proteinase K-treated small RNA preparations suggests that these proteins possess three essential chemical properties that allow their co-purification and protection during glycoRNA isolation^3^: i) hydrophilic phase separation during organic phenol-chloroform extraction, ii) binding to silica columns, and iii) relative resistance to proteinase K treatment under non-denaturing conditions. Significantly, most proteins identified here are known either to be glycosylated or to possess RNA-binding capabilities, often comprising both features (Supplementary Table 1, 2). Whereas 10 proteins were commonly identified between 3T3 and HeLa cells (Supplementary Figure 2e, f), the identification of individual hits, unique for either 3T3 or HeLa cells, is likely related to cell type- and species-specific expression and modification patterns. Here, we used LAMP1 as a representative glycoprotein from the group of commonly identified proteins to study the possibility of glycoprotein co-purification during glycoRNA isolation. We demonstrated that purified sol-hLAMP1 at least partially localized to the aqueous phase during TRIzol extraction (Figure 5b), which is likely mediated by its extensive decoration with glycans^25,26^, which can significantly enhance hydrophilicity^2^. Indeed, it is established that, along with association to RNA, glycan-modification can disturb “canonical” phase separation^30^. Exemplarily, contamination of RNA and DNA samples, extracted from blood or tissue, with the glycosaminoglycan heparin is a known phenomenon^31,32^. Besides, the cytosolic tail of LAMP1 was recently reported to bind RNA^33^, and the intrinsically disordered region (IDR) within the hinge region of LAMP1 might exert non-classical modes of RNA-binding as seen in condensates^34^, membrane-less organelles that are known to incorporate RNA^35^. These aspects might also promote the co-purification of glycoproteins like LAMP1 during the purification of glycoRNA following the published protocol^3^.

Recently, Kim et al. showed that the presence of RNA and higher concentrations of alcohol are important for the co-purification of unknown N-glycosylated molecules during glycoRNA isolation^17^, which we were able to reproduce (Figure 3c). In our proteomics approach (Figure 4), we separated large and small RNAs based on the alcohol concentration during column loading (using high alcohol concentrations for small RNA separation), and we observed that sol-hLAMP1 was recovered from the column purification dependent on the presence of RNA or higher concentrations of alcohol (Figure 5). Thus, we conclude that the findings by Kim and co-workers also apply to glycoproteins.

Regarding resistance to proteinase K, it has been shown that extensive glycosylation can protect from proteolytic cleavage either by shielding cleavage sites or via electrostatic repulsion^2^. For example, O-glycosylated fragments of mucins were found to be largely resistant to proteinase K cleavage, a property that has been exploited to isolate these from other proteins that are more susceptible to digestion^36,37^. In the case of LAMP1, relative protection from proteinase K is expected, given that the glyco-matrix of LAMP1^25,26^ helps to protect the endosome from autolysis^38,39^. Although proteinase K is active across a wide range of buffers and pH^40^, its activity can be stimulated upon the addition of denaturing and reducing agents, which generate access to hidden cleavage sites in heavily structured target proteins^19^. We observed that the use of a buffer system with denaturing and reducing activity leads to the removal of glycoprotein contaminants from glycoRNA preparations (Figure 2). As high nuclease resistance of glycosylated molecules in RNA preparations was reported by other studies before^12,17,41^, we hypothesize that the co-purification of glycoproteins during glycoRNA isolation is a common phenomenon. Future investigations of glycoRNA may adapt the herein described proteinase K treatment under denaturing conditions, which may allow for the fine-tuning of the established method towards higher glycoRNA purity. On the other hand, RNA isolation under conditions with no or mild proteinase K digestion may enrich for glycosylated and/or RNA-binding proteins.

In summary, we here describe the co-purification of glycoproteins during the preparation of glycoRNA following the recently described method^3^. We hypothesize that the glycosylation status and RNA-binding properties of the proteins identified herein mediate both their co-purification and protection during isolation. The implementation of a proteinase K digestion step under denaturing conditions may allow for further removal of glyco-molecules of non-RNA origin, enabling the investigation of further structural and functional aspects of glycoRNA.

## Supporting information

Supplementary data file (MaxQuant)

## Acknowledgements

The authors would like to thank Lisa Mainieri, Vincent Kramer, and Dr Jens Dorna for help with method development, and Dr Francisco Venegas Solis, Prof. PhD Johannes Graumann, and Dr Witold Szymanski for excellent chemical expertise. Moreover, we would like to thank Dr Nadiia Pozhydaieva for her help with the proteomics data analysis.

This work was supported by Deutsche Forschungsgemeinschaft (DFG, German Research Foundation) Project-ID 369799452 – TRR237 - A02 to SB. This work has received funding from the European Research Council under the European Union’s Horizon 2020 research and innovation programme (ERC-2023-STG grant 101114948, NAD-ART to K.H.), and from the Max Planck Society (Max Planck Research Group Leader funding to KH).

## Author contributions

**NBK**: conceptualization; investigation; methodology; visualization; writing – original draft preparation. **NYD**: conceptualization; data curation; formal analysis; investigation; methodology; visualization. **TG**: data curation; formal analysis; investigation; resources. **KH**: conceptualization; funding acquisition; methodology; project administration; supervision. **AK**: conceptualization; methodology; supervision; **SB**: conceptualization; funding acquisition; methodology; project administration; supervision; writing – original draft preparation. **All authors**: writing – review & editing.

---

**Supplementary Figure 1, related to Figure 2:**
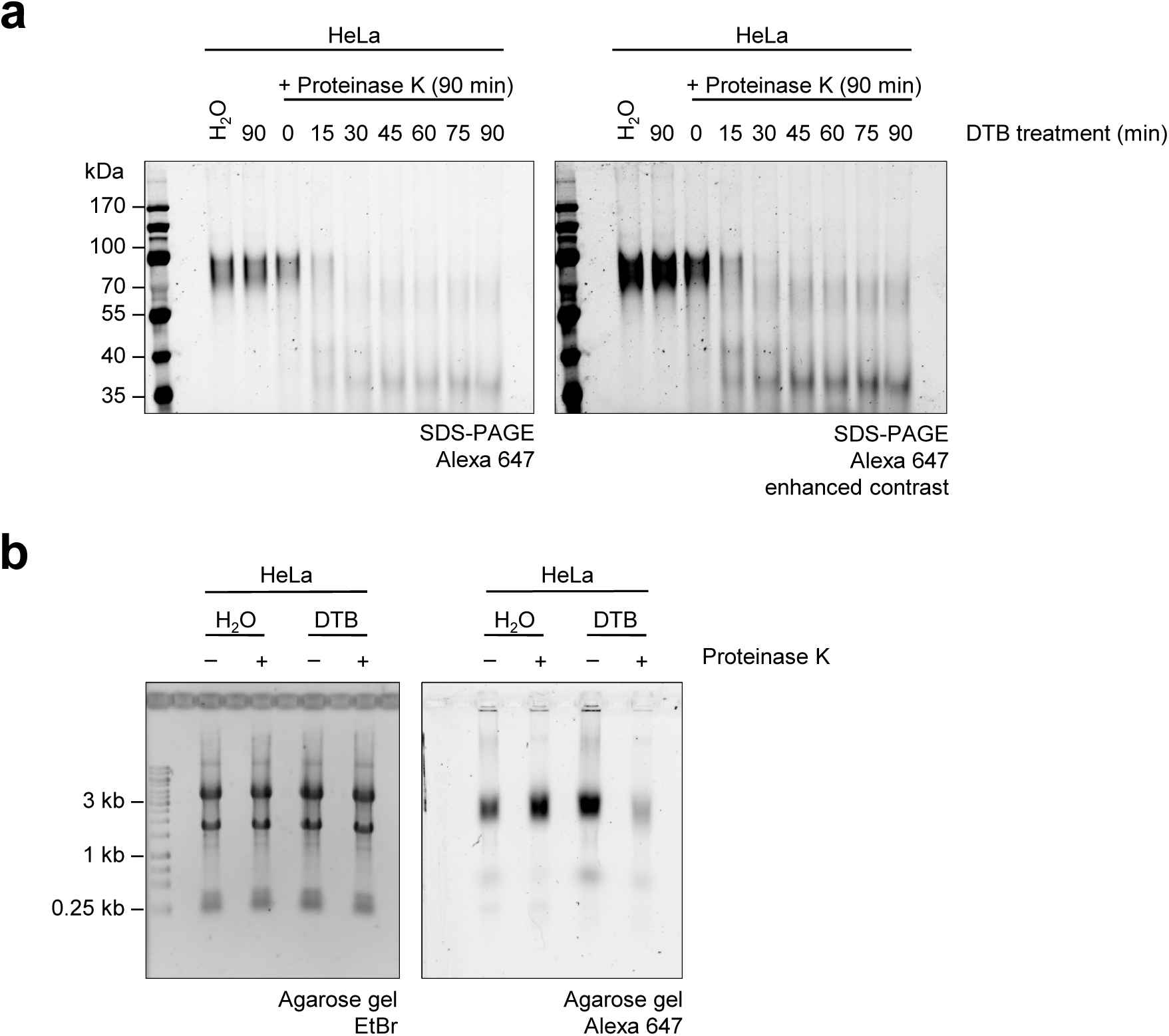
Proteinase K treatment of HeLa RNA in denaturing buffer: Experiments in (a) and (b) were conducted as in Figure 2b and 2d using DBCO-AF647-clicked RNA from metabolically labelled HeLa cells.

**Supplementary Figure 2, related to Figure 4:**
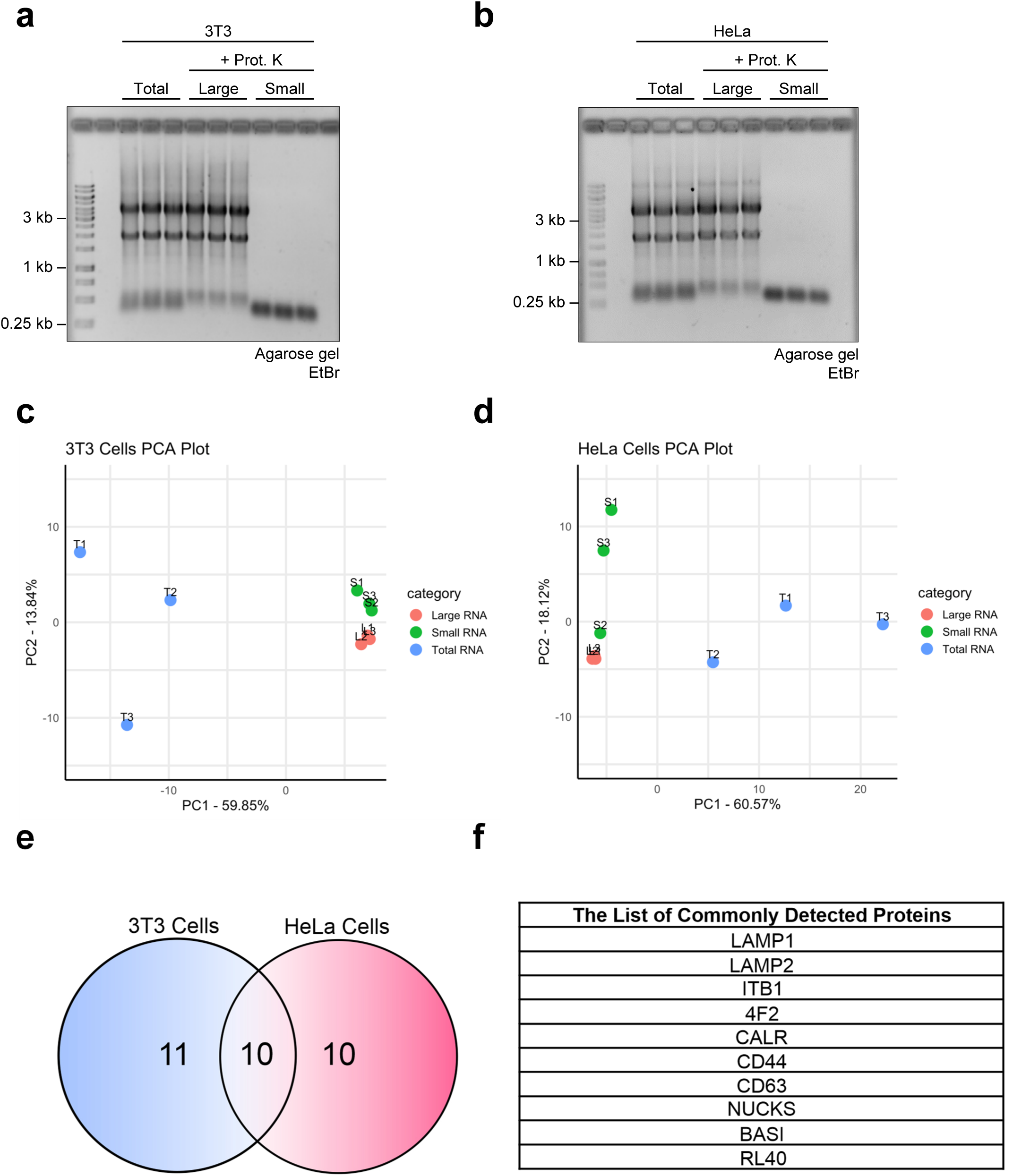
RNA samples for proteomics analysis, PCA results, and commonly detected proteins. (a) Agarose gel with EtBr staining of the three sample groups (total RNA, large RNA fraction, small RNA fraction) extracted and purified from 3T3 cells. (b) Agarose gel as in (a) but with samples obtained from HeLa cells. (c) Principal Component Analysis (PCA) of sample label-free quantification (LFQ) intensities to evaluate the reproducibility of triplicates. The score plot illustrates the distribution of individual samples along the first two principal components (PC1 and PC2). PCA of samples from 3T3 cells showing the close clustering of replicates from the same sample set. (d) PCA of samples from HeLa cells showing the close clustering of replicates from the same sample set. (e) Venn diagram illustrating the number of individually detected proteins as well as commonly detected proteins between 3T3 and HeLa cells. (f) Table displaying the list of commonly detected proteins.

**Supplementary Table 1:** Proteins with an average unique peptide count of ≥2 identified in small RNA fraction samples extracted from 3T3 cells with information on molecular weight, glycosylation, and RNA-binding properties.

| **Protein IDs**<br>Uniprot accession number | **aa / MW (kDa)** | **Glycosylation sites reported on UniProt**<br>including inferred sites based on sequence analysis | **Known interaction with RNA**<br>including occurrences in high-throughput analyses |
| --- | --- | --- | --- |
| **ANXA2**<br>P07356 | 339 / 38.7 | None, but probably glycosylated in humans (PMID: 1445299) | No |
| **DESP**<br>E9Q557 | 2883 / 332.9 | None | No |
| **PLAK**<br>Q02257 | 745 / 81.8 | 1 O-linked (sequence analysis, PMID: 12847106) | Yes (27281784, 29339797) |
| **LAMP1**<br>P11438 | 406 / 43.9 | 18 N-linked (PMID: 27663661, 16170054, 19349973, 19656770, 16944957) | Yes (PMID: 25772617) |
| **LAMP2**<br>P17047 | 415 / 45.7 | 16 N-linked (PMID: 2766366) | Yes (PMID: 23291500) |
| **CD34**<br>Q64314 | 382 / 41.0 | 7 N-linked (PMID: 19656770) | No |
| **CD44**<br>P15379 | 778 / 85.6 | 10 N-linked (sequence analysis)<br>1 O-linked (by similarity, see human CD44) | No |
| **4F2**<br>P10852 | 526 / 58.3 | 8 N-linked (PMID: 19349973, 19656770) | Yes (PMID: 21266579) |
| **LPR1**<br>Q91ZX7 | 4545 / 504.7 | 51 N-linked (PMID: 19656770, 19349973) | No |
| **LAMB1**<br>P02469 | 1786 / 197.1 | 12 N-linked (PMID: 19349973) | No |
| **LAMC1**<br>P02468 | 1607 / 177.3 | 14 N-linked (PMID: 19349973) | No |
| **BASI**<br>P18572 | 389 / 42.4 | 3 N-linked (PMID: 19349973, 19656770) | No |
| **EIF3A**<br>P23116 | 1344 / 161.9 | O-glycosylated (PMID: 34887587) | Yes (PMID: 17581632) |
| **MPRI**<br>Q07113 | 2483 / 273.8 | 20 N-linked (PMID: 19349973, 19656770) | No |
| **ITB1**<br>P09055 | 798 / 88.2 | 12 N-linked (PMID: 19656770) | No |
| **PGBM**<br>Q05793 | 3707 / 398.3 | 4 O-linked (sequence analysis + by similarity)<br>10 N-linked (PMID: 19656770, 19349973) | No |
| **FINC**<br>P11276 | 2477 / 272.5 | 8 N-linked (PMID: 17330941, 19656770, 16944957) | No |
| **CALR**<br>P14211 | 416 / 48.0 | 1 N-linked (see human CALR)<br>binds to all monoglycosylated proteins in the Golgi apparatus (PMID: 20880849, 21652723) | Yes (in humans; PMID: 12242300) |
| **RL40**<br>P62984 | 128 / 14.7 | None | Part of the 60S ribosomal subunit (PMID: 19754430, 36517592) |
| **CD63**<br>P41731 | 238 / 25.8 | 4 N-linked (PMID: 19349973) | No |
| **NUCKS**<br>Q80XU3 | 234 / 26.3 | None | Yes (in humans: PMID: 22681889) |

**Supplementary Table 2:** Proteins with an average unique peptide count of ≥2 identified in small RNA fraction samples extracted from HeLa cells with information on molecular weight, glycosylation, and RNA-binding properties.

| **Protein IDs**<br>Uniprot accession number | **aa / MW (kDa)** | **Glycosylation sites reported on UniProt**<br>including inferred sites based on sequence analysis | **Known interaction with RNA**<br>including occurrences in high-throughput analyses |
| --- | --- | --- | --- |
| **NUCKS**<br>Q9H1E3 | 243 / 27.3 | None | Yes (PMID: 22681889) |
| **PTMA**<br>P06454 | 111 / 12.2 | None | Yes (PMID: 2452757, 2479575, 9326492) |
| **GLYG**<br>P46976 | 350 / 39.4 | 1 O-linked (PMID: 22160680, 30356213) | No |
| **RL40**<br>P62987 | 128 / 14.7 | None | Is a component of the 60S ribosomal subunit (PMID: 23169626, 23636399, 32669547, 39048817, 39103523) |
| **L1CAM**<br>P32004 | 1257 / 140.0 | 20 N-linked (PMID: 16335952, 19159218) | No |
| **NUCL**<br>P19338 | 710 / 76.6 | A surface-located form of NUCL is glycosylated (PMID: 21575138, 15823039, 19023103, 19026635) | Yes (PMID: 8321232, 16213212) |
| **4F2**<br>P08195 | 630 / 68.0 | 4 N-linked (PMID: 12754519, 19159218, 19349973, 30867591, 33298890, 33758168, 34880232, 35352032, 16335952, 19139490) | Yes (PMID: 21266579, 22658674) |
| **CD63**<br>P08962 | 238 / 25.6 | 3 N-linked (PMID: 12754519, 19159218) | No |
| **BST2**<br>Q10589 | 180 / 19.8 | 2 N-linked (PMID: 19159218, 19349973, 19737401, 19879838) | Yes (PMID: 22658674) |
| **ITB1**<br>P05556 | 798 / 88.4 | 12 N-linked (PMID: 19159218, 22451694, 33962943, 16335952, 19349973) | No |
| **BASI**<br>P35613 | 385 / 42.2 | 3 N-linked (PMID:<br>12754519, 19159218,<br>19349973) | No |
| **CALR**<br>P27797 | 417 / 48.1 | 1 N-linked (PMID:<br>19159218)<br>binds to all<br>monoglycosylated<br>proteins in the ER<br>(PMID: 7876246) | Yes (PMID: 14726956,<br>22658674) |
| **LAMP1**<br>P11279 | 417 / 44.9 | 18 N-linked (PMID:<br>3143719, 12754519,<br>19159218, 16335952)<br>6 O-linked (PMID:<br>8323299) | Yes (PMID: 25772617) |
| **CD44**<br>P16070 | 742 / 81.5 | 9 N-linked (PMID:<br>12883358, 16335952,<br>19159218, 22171320)<br>1 O-linked (PMID:<br>25326458, 32337544,<br>36213313, 37453717) | No |
| **LAMP2**<br>P13473 | 410 / 45.0 | 16 N-linked (PMID:<br>2243102, 2912382,<br>8323299, 12754519,<br>16335952, 19159218)<br>10 O-linked (PMID:<br>8323299) | Yes (PMID: 23291500) |
| **TGON2**<br>O43493 | 437 / 45.9 | 9 N-linked (sequence<br>analysis) | No |
| **MUC16**<br>Q8WXI7 | 14507 / >1500 | 102 N-linked, heavily<br>O-glycosylated (PMID:<br>12734200), | No |
| **CALX**<br>P27824 | 592 / 67.6 | None, but functions as<br>a chaperone during<br>glycosylation quality<br>control in the ER<br>(PMID: 8203019,<br>7736594) | Yes (PMID: 22658674,<br>22681889) |
| **AN32A**<br>P39687 | 249 / 28.6 | None | Yes (PMID: 22681889) |
| **NPM**<br>P06748 | 294 / 32.6 | None | Yes (PMID: 12058066,<br>24106084) |

